# Microneedle-mediated intratumoral delivery of anti-CTLA-4 promotes cDC1-dependent eradication of oral squamous cell carcinoma with limited irAEs

**DOI:** 10.1101/2021.12.07.471658

**Authors:** Mara Gilardi, Robert Saddawi-Konefka, Victoria H. Wu, Miguel Angel Lopez-Ramirez, Zhiyong Wang, Fernando Soto, Dana J. Steffen, Marco Proietto, Zbigniew Mikulski, Haruka Miki, Andrew Sharabi, Daniel Kupor, Ricardo Rueda, Daniel P. Hollern, Joseph Wang, J. Silvio Gutkind

**Affiliations:** Moores Cancer Center, University of California San Diego, 3855 Health Sciences Drive, La Jolla, CA, 92093, USA; Division of Otolaryngology-Head and Neck Surgery, Department of Surgery, University of California San Diego, La Jolla, CA, 92093, USA; Department of Pharmacology, University of California San Diego, La Jolla, CA, 92093, USA; Department of Nanoengineering, University of California San Diego, La Jolla, CA, 92093, USA; Section of Cell and Developmental Biology, University of California San Diego, La Jolla, CA, 92093, USA; Microscopy Core Facility, La Jolla Institute for Immunology, La Jolla, CA, 92093, USA; Center of Autoimmunity and Inflammation, La Jolla Institute for Immunology, La Jolla, CA, 92093, USA; Department of Radiation Medicine and Applied Sciences, University of California San Diego, La Jolla, CA, 92093, USA; Salk cancer center, La Jolla, CA, 92037, USA; Nomis cancer center for immunology and microbial pathogenesis, La Jolla, CA, 92037, USA

**Keywords:** Head and neck cancer, microneedle, immunotherapy, intratumoral delivery, cDC1

## Abstract

Head and neck squamous cell carcinoma (HNSCC) ranks 6th in cancer incidence worldwide and has a five-year survival rate of only 63%. Immunotherapies – principally immune checkpoint inhibitors (ICI), such as anti-PD-1 and anti-CTLA-4 antibodies that restore endogenous antitumor T-cell immunity – offer the greatest promise for HNSCC treatment. Anti-PD-1 has been recently approved for first line treatment of recurrent and metastatic HNSCC; however less than 20% of patients show clinical benefit and durable responses. In addition, the clinical application of ICI has been limited by immune-related adverse events (irAEs) consequent to compromised peripheral immune tolerance. Although irAEs are often reversible, they can become severe, prompting premature therapy termination or becoming life-threatening. To address the irAEs inherent to systemic ICI therapy, we developed a novel, local delivery strategy based upon an array of soluble microneedles (MN). Using our recently reported syngeneic, tobacco-signature murine HNSCC model, we found that both systemic and local-MN anti-CTLA-4 therapy lead to >90% tumor response, which is dependent on CD8 T cells and conventional dendritic cell type 1 (cDC1). However, local-MN delivery limited the distribution of anti-CTLA-4 antibody from areas distal to draining lymphatic basins. Employing Foxp3-*GFP*^DTR^ transgenic mice to interrogate irAEs *in vivo*, we found that local-MN delivery of anti-CTLA-4 protects animals from irAEs observed with systemic therapy. Taken together, our findings support the exploration of MN-intratumoral ICI delivery as a viable strategy for HNSCC treatment with reduced irAEs, and the opportunity to target cDC1s as part of multimodal treatment options to boost ICI therapy.

## Introduction

Head and neck squamous cell carcinoma (HNSCC) is an often lethal disease, accounting for 62,000 new cases and 13,000 deaths annually in the United States alone (1). Despite advances in curative-intent therapies over the past three decades, long-term toxicities remain unacceptably morbid, and the overall mortality rate remains quite high and recurrences occur in as many as half of all initial responders (1,2). Recently, immune oncology (IO) therapies – namely, immune checkpoint inhibitors (ICI) anti-PD-1/PD-L1 (αPD-1/αPD-L1) and anti-CTLA-4 (αCTLA-4) – have provided new and effective clinical treatment strategies for cancer patients. ICI therapy restores exhausted CD8 T cell antitumor immunity and offers great promise. Based on results in large phase III clinical trials, αPD-1 is now approved for the management of recurrent/metastatic HNSCC as both first- and second-line therapy (3,4). However, less than 20% of HNSCC patients show clinical benefit and durable responses to αPD-1 (3,4). αCTLA-4 trials in recurrent/metastatic HNSCC have fared less well, but emerging neoadjuvant αPD-1 monotherapy and combination αPD-1/αCTLA-4 trials show synergism that can be exploited therapeutically (5,6).

Of interest, excitement over clinical responses to ICI therapy have been tempered by the reality of immune-related adverse events (irAEs), which occur in roughly 54% of cases after αCTLA-4 monotherapy with dreaded grade 3/4 irAEs occurring in 10-20% or 55-60% of cases after αPD-1 or combination ICI treatments, respectively (7,8). Although irAEs are often reversible, they can become severe, at best prompting premature termination of therapy or at worst becoming life-threatening, with irAE-related fatalities occurring in 0.3-1.3% of cases (8). The goal of ICI therapy, particularly αCTLA-4 blockade, is to reduce the threshold for CD8 T cell immune activation and augment responses against tumor antigens, thereby mediating tumor rejection. As ICI therapies are delivered systemically to patients, the threshold for immune tolerance is likewise lowered peripherally, leading to irAEs (8). Dose-limiting irAEs represent an immediate clinical obstacle, highlighting the need to develop effective and less toxic therapeutic strategies and delivery options.

A defining feature of most HNSCCs as well as oral premalignant lesions (OPL) is the superficial and mucosal localization of the disease. Unlike many other cancers, most HNSCC lesions can be readily visualized and accessed, providing opportunities for intratumoral (IT) drug delivery. Using syngeneic HNSCC models, we recently demonstrated that IT delivered αCTLA-4 achieves similar therapeutic responses to systemic delivered (9). However, reliable IT delivery is difficult to achieve. Instead, dissolvable microneedle (MN) devices represent an attractive alternative to hypodermic needles as they offer painless and minimally invasive therapeutic payload delivery within a few microns of the target lesion, require minimal training to be self-applied, leave no sharp biohazardous waste, and have long shelf life (10).

Here, we report an engineered dissolvable and biodegradable *in-situ* MN patch for the intratumoral delivery of αCTLA-4 antibody towards overcoming the immune checkpoint inhibitor resistance in HNSCC. We found that this localized treatment achieves durable responses with more localized distribution over systemic delivery. We also describe the mechanism of tumor rejection downstream from αCTLA-4 MN delivered treatment as both CD8 T-cell- and conventional type I dendritic cell (cDC1)-dependent. Importantly, the MN delivery system protects hosts from dose-limiting irAEs, which are otherwise prohibitive. The use of MN arrays may provide a promising and novel modality for the IT delivery of immunotherapies in HNSCC.

## Materials and Methods

### Cell lines and tissue culture

The 4MOSC1 syngeneic mouse HNSCC cells harboring a human tobacco-related mutanome and genomic landscape were developed and described for the use in immunotherapy studies in our prior report (9). MOC1 syngeneic mouse HNSCC cells derived from DMBA-induced oral tumors were generously provided by Dr. R. Uppaluri (11).

### Reagents

Agarose, Polyvinylpyrrolidone (PVP) Mw∼360K, Rhodamine 6G (Rh6G), Fluorescein 5(6)-isothiocyanate (FITC) were purchased from Sigma Aldrich. IgG-AlexaFluor555 from Abcam. Dow SYLGARD® 184 silicone encapsulant from Ellsworth Adhesives. CTLA-4 antibody (clone 9H10, catalog #BP0131), isotype antibody (catalog # BE0091), and CD8 depletion antibody (Clone YTS 169.4, catalog #BE0117) were obtained from Bio X Cell (West Lebanon, NH, USA). Fluorochrome-conjugated antibodies were purchased from BioLegend and BD Biosciences.

### Negative PDMS MN mold fabrication

A master MN mold made of acrylate resin was attached to a Crystal Clear Borosillicate Glass petri dish with a double-sided tape. A PDMS solution with a ratio of 8.6/1.4 (base/curing agent) was prepared and casted onto the MN mold. Furthermore, the MN mold was placed within a sealed desiccator at 23 in Hg for 5 min. PDMS was set under room temperature for 1 hour and 30 min, afterwards placed within an oven for an additional 30 min at 85°C to harden/complete the curing process. The negative MN mold was detached from the substrate and the size adjusted with the use of a blade cut. Additionally, MN molds were washed by triplicate with hand soap and rinsed with water, ultrasonicated for 15 min, cleaned with 2-propanol and placed in the oven at 75°C for 15 min. Sterilized molds were stored in a sealed contained at RT until use.

### Fabrication of αCTLA-4 MN patches

A volume of 60µL of a 10% w/v PVP aqueous solution fabricated at pH 7.4 was casted onto negative PDMS MN molds and placed in a sealed desiccator under vacuum (5 min at 23 in Hg). Subsequently, MN molds were carefully transferred out of the desiccator, and bubbles generated and trapped within each MN negative feature were removed. Moreover, repetitive additions of PVP were implemented, until reaching a total volume of 240µL. 100µg of αCTLA-4, 100 µg of IgG (control), 50 µg of IgG Alexa Fluor-555 (imaging), 50 µg of FITC (imaging) or 50 µg Rh6G (imaging) was casted to the mold and was let to dry for 24 hours. After the drying process, a 7 × 7 mm 3M scotch tape was applied onto the polymeric film and the MN patch was peeled off from the silicone negative mold. After demolding, the MN patches were stored in a sealed container at 4°C until use. Control/blank MN patches were fabricated following the same preparation procedure in the absence of αCTLA-4 or IgG.

### *In vivo* mouse experiments and analysis

All the animal studies using HNSCC tumor xenografts studies were approved by the Institutional Animal Care and Use Committee (IACUC) of University of California, San Diego, with protocol ASP # S15195. Female C57Bl/6 mice (4–6 weeks of age and weighing 16–18 g) were purchased from Charles River Laboratories (Worcester, MA, USA). For Foxp3^DTR^ mice, we use C57BL/6-Tg(Foxp3-DTR/EGFP mice (stock # 016958); from JAX in C57BL/6 background) (6–8 weeks of age and weighing 18–22 g). To deplete Tregs, mice were injected IP with 500 ng of diphtheria toxin (DT; Sigma-Aldrich), diluted in PBS, as we recently described (9). For Batf3 KO mice, we use B6.129S(C)-Batf3^tm1Kmm^/J mice (stock # 013755) from JAX in C57BL/6 background) (4–6 weeks of age and weighing 18–22 g). Tumor studies and histopathology analysis were performed as we recently described (9).

### Immunofluorescence and image quantification

Briefly, after the treatment with ICB, tissues (tongue, cervical lymph nodes, and spleen) were harvested, fixed, and paraffin embedded. Tissues were stained for Cytokeratin 5 (CK5, Fitzgerald, 20R-CP003) (1:500), CD8 (Abcam, ab22378) (1:400), anti-Syrian hamster IgG (Abcam, ab180117) and CD11c (Abcam, ab219799) antibodies. Secondary antibodies were used to reveal the specific marker signal in multiplex immune fluorescence (IF). Hematoxylin and eosin (H&E) staining was performed on tissue sections for histopathology analysis. Zeiss 780 confocal microscope was used for fluorescence imaging. Quantification of immune infiltration was performed using QuPath, an open-source software for digital pathology image analysis (12). Zeiss Axioscan was used to scan the H&E-stained sections. For the quantification, at least three regions of interest (ROI) were selected for each condition and the percentage of positive cells for each marker was calculated as we previously reported (13).

### Tumor infiltrating lymphocyte isolation and flow cytometry

Tumors and lymph nodes were dissected, minced, and resuspended in complete media (DMEM with 10% FBS and 1% antibiotics) supplemented with Collagenase-D (1 mg/mL; Roche) and incubated at 37°C for 30 min with shaking to form a single-cell suspension. Tissue suspensions were washed with fresh media and passed through a 100-µm strainer. Samples were washed with PBS and immediately processed for live/dead cell discrimination using BD Horizon™ Fixable Viability Stain 510. Cell surface staining was done for 30 min at 4°C with the antibodies as previously described (9,14). Markers for cDC1 populations were characterized as previously described (15). All flow cytometry was run using BD LSR Fortessa and analyzed using FlowJo. Tumor infiltrating lymphocyte (TIL) count was determined using CountBright™ Absolute Counting Beads (ThermoFisher Scientific). Immune cells were identified as previously described (9).

### Histology and inflammatory cell analysis

Tissues (skin, eye lid, liver, lung, and colon) were harvested, fixed, and paraffin embedded, and H&E stained. Images were acquired by Axioscan (Zeiss). Quantification of immune infiltration has performed using QuPath, an open-source software for digital pathology image analysis (12). For the quantification, the whole tissue section was scanned, annotated and analyzed for each condition with the pixel classifier training function. On the H&E slide, after the tissue annotation defining the ROI for the quantification, the cell detection was performed by QuPath. The pixel classifier for inflammation was trained based on immune cell infiltration detection for each ROI. The pixel classifier was trained differently for each type of tissue given the tissue heterogeneity and staining background. The percentage of positive cells for each marker was calculated respect of the total number of cells, as we previously reported (9,13).

## Results

### Fabrication and characterization of micro-needle (MN) patches for the delivery of αCTLA-4

To effectively use MN patches to eradicate the growth of HNSCC in the tongue of mice, their design and fabrication were performed to fit the appropriate dimensions of the target area. A schematic illustration shows the oral dissolvable MN application on 4MOSC1 HNSCC tumors (**Fig. 1A**), the width of each MN patch measured 1cm. The MN platform is comprised of both, the therapeutic (αCTLA-4) IO agent and the polyvinylpyrrolidone (PVP) polymeric matrix. The hydrophilicity and biocompatible properties of PVP has allowed its widespread use in the biomedical industry and clinical setting, providing low cytotoxicity and biodegradability (16). Briefly, MNs were fabricated by the micromolding approach (17) (**Supplemental Fig. 1**). A negative poly(dimethylsiloxane) (PDMS) MN mold was employed as a template, where its cavities were loaded with the IO agent embedded PVP and followed by drying the patch overnight. The finalized replica MN patch was then transferred to a medical adhesive support and stored at 4°C in a sealed container prior use. Our polymeric MN patch was prepared under gentle and soft experimental conditions, therefore avoiding the use of harsh organic solvents or elevated temperatures capable of denaturation of the entrapped IO agent. These tailored-made PVP/αCTLA-4 dissolvable MNs were designed to dissolve upon contact with tumoral tissue and biological fluids present in the oral cavity (saliva). The MN design, materials and device fabrication were chosen to guardedly enclose the IO agent while preserving antigenicity, leaving behind only safe soluble products. As the polymer matrix responsible for the rigid structure of each MN dissolves, the release of the IO agent to the target area proceeds within minutes (17) (**Supplemental Fig. 1**). Specifically, complete local delivery was confirmed following MN detachment from the underlying support, and kinetic release was achieved within 30 minutes after the treatment (**Supplemental Fig. 1**).

**Fig. 1.**
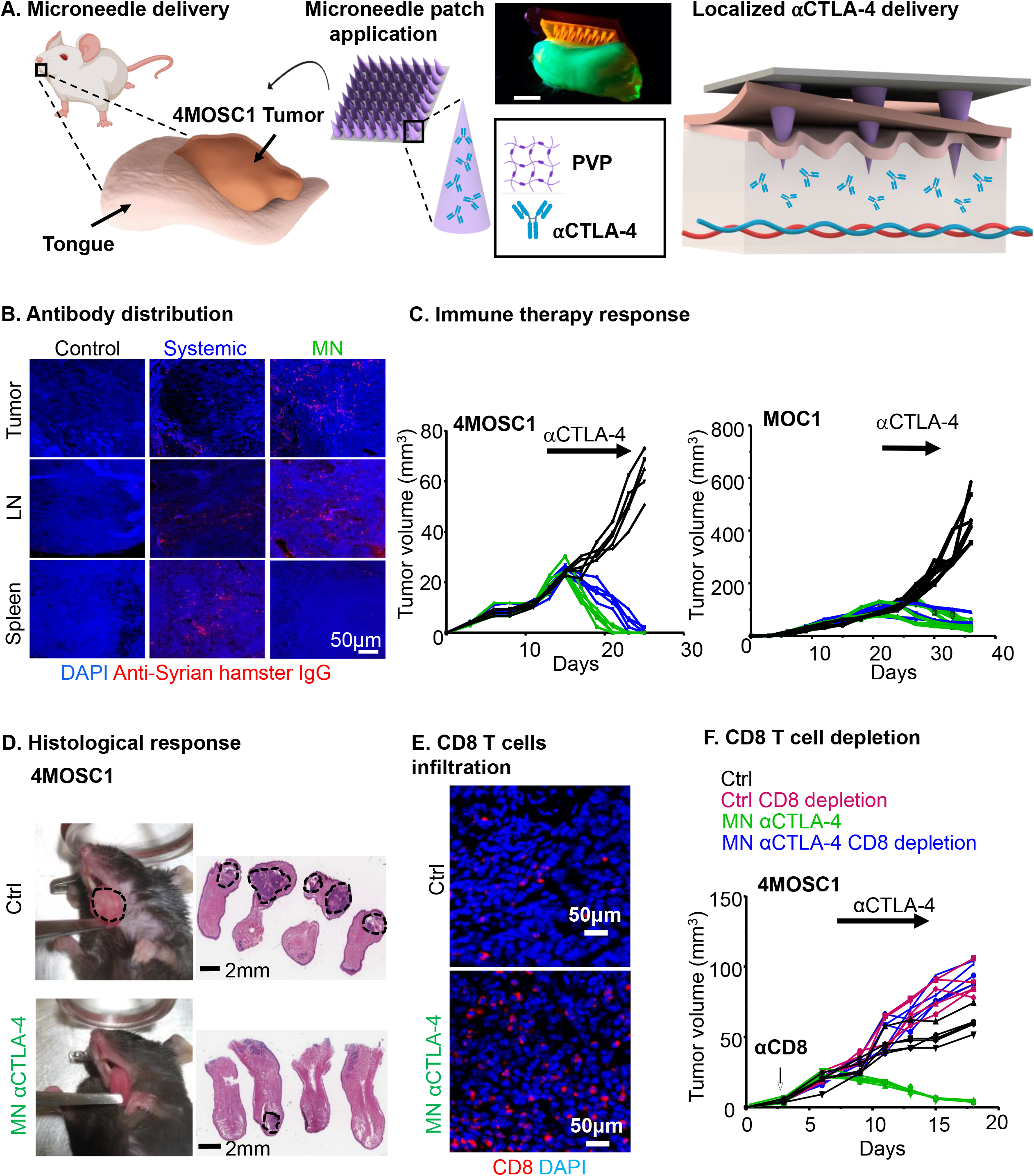
Microneedle patches as a localized αCTLA-4 delivery system. **A)** Schematic of a dissolvable MN application on 4MOSC1 tumors in the tongue of mice for the delivery and release of an IO agent (αCTLA-4) to treat HNSCC. Digital photograph of the application of a dye supplemented (Rh6G) MN patch onto a fluorescent synthetic FITC hydrogel mouse tongue. Scale bar, 3mm. **B-D)** C57Bl/6 mice were implanted with 1 × 10^6^ 4MOSC1 cells into the tongue. After the tumors reached ∼30 mm^3^, mice were treated with PVP patches (MN, control), or αCTLA-4 systemic (5 mg/kg) or by MN (0.1 mg) delivery. **B**). Shown is the immunofluorescent staining of the distribution of anti-Syrian hamster IgG (isotype for αCTLA-4 antibody) in mice with 4MOSC1 tumors treated by negative MN (control), αCTLA-4 s (IP, systemic) or αCTLA-4 MN. Staining for anti-hamster IgG (red) showed the localization of αCTLA-4 antibody in the tongue, lymph nodes, and spleen of treated mice. DAPI staining for nuclei is shown in blue (representative from 3 independent experiments with each n = 3 mice per group). **C**) Individual growth curves of 4MOSC1 and MOC1 tumor-bearing mice. Primary tumor growths are shown (n = 5 mice per group) using control (black), αCTLA-4 IP (blue), or αCTLA-4 MN (green). **D**) Histological responses. Left panel, representative pictures of tongues from isotype control and αCTLA-4 MN groups. Middle panel, representative H&E staining of histological tissue sections from mouse tongues from isotype control and αCTLA-4 MN groups. Sale bars represent 2mm. **E)** Immunofluorescence staining showing increased CD8 T-cells recruitment in the tumor due to MN αCTLA-4 treatment. Scale bars represent 50μm. **F)** Dependency of MN αCTLA-4 on CD8 T-cells. C57Bl/6 mice were treated with CD8 T-cell depleting antibody daily for 3 days before tumor implantation and then once a week after. HNSCC tumor bearing mice (as above) were treated with negative PVP MN (control) or αCTLA-4 MN (twice a week) (n = 5 per group). Individual growth curves of 4MOSC1 tumor-bearing mice are shown.

### MN local delivery of αCTLA-4 exerts therapeutic potential in HNSCC syngeneic model

We have previously observed single-agent response to αCTLA-4 in our novel murine, tobacco-signature HNSCC model (4MOSC model (9)). To determine the efficacy of local versus systemic αCTLA-4 delivery, we developed a first generation of polymeric/αCTLA-4 MN patch, as described above. To explore the distribution of therapeutic antibody in the host following treatment, we performed immunofluorescent analysis of tumor, lymph node and splenic compartments after 24h from both systemic and local-MN delivery. Tumor lesions were treated when the volume reached 20-30 mm^3^ respectively with, 5mg/kg (systemic, approximately 0.1 mg per mouse) and 0.1mg (for each MN patch) of αCTLA-4 antibody (**Fig. 1B**). Indeed, dose response experiments supported that 0.1mg of αCTLA-4 represents a minimum effective dose for *in vivo* treatment (**Supplemental Fig. 2**). Opposite from the pattern observed 24 hours after after systemic delivery, local-MN delivery resulted in a greater and localized distribution of αCTLA-4 antibody in the tumor and draining lymph node compartments with restricted distribution more distally to the spleen, as a representative peripheral tissue. This local application of αCTLA-4 loaded-MN patches onto established 4MOSC1 tumors elicited a robust antitumor effect, with most tumor-bearing mice exhibiting complete responses (CR) as observed with systemic application (**Fig. 1C & 1D**). Similar results were obtained in different syngeneic mouse model, MOC1, grown in the flank of mice (**Fig. 1C**)). We next addressed whether local application of αCTLA-4 could lead to a significant increase in CD8 T-cell infiltration. As expected, we observed an increase in CD8 T-cells infiltrating into tumors after local αCTLA-4 therapy (9) (**Fig. 1E**). To determine the necessity of CD8 T-cells in the tumor response to MN delivered αCTLA-4, we treated tumor-bearing animals with a depleting CD8 antibody and observed that all CD8-depleted animals fail to respond to local αCTLA-4 therapy (**Fig. 1F**).

### αCTLA-4 requires host conventional type I dendritic cells (cDC1)

Noting that local αCTLA-4 delivery via the MN leads to a preferentially proximal redistribution of therapeutic antibody and that treatment-response requires CD8 T-cells, we examined conventional type I dendritic cell (DC) populations (cDC1) in both the tumor and draining lymph nodes (DLN) (**Fig. 2A**). Immunofluorescence shows a robust accumulation of dendritic cells (CD11c+) into the tumor and DLN compared to control. Next, we sought to examine for the presence of conventional type I DCs (cDC1). cDC1s with the ability to efficiently cross-present cell-associated antigens are recognized as key effectors in priming antitumor cytotoxic lymphocytes (18,19). To examine cDC1s in our system, we performed flow cytometric analysis to detect the presence of either tumor-associated cDC1s (CD11c^+^CD103^+^CD11b^−^) or cervical lymph node-associated cDC1s (CD11c^+^CD8a^+^CD11b^−^) after either systemic or local αCTLA-4 therapy (**Fig. 2B**, and **Supplemental Fig. 3**). We found a significant increase in intratumoral and lymph node populations of cDC1s after systemic and MN delivery of αCTLA-4 (**Fig. 2B**). Concomitant with this, we observed a significant increase of CD8 T cells in the LN, suggesting that the DC priming event is associated with increased T cell responses (**Supplemental Fig. 4)**. To compare the phenotype of cDC1s that accumulate in the lymph nodes we measured the expression of several key functional receptors or cytokines with known roles in T cell priming and recruitment. Among them, the expression of CD40 and CCR7, as well as IL-12 secretion, were found to be significantly increased in cDC1 in lymph nodes after systemic and locally delivered αCTLA-4 treatment (**Fig. 2C** and **2D**).

**Fig. 2.**
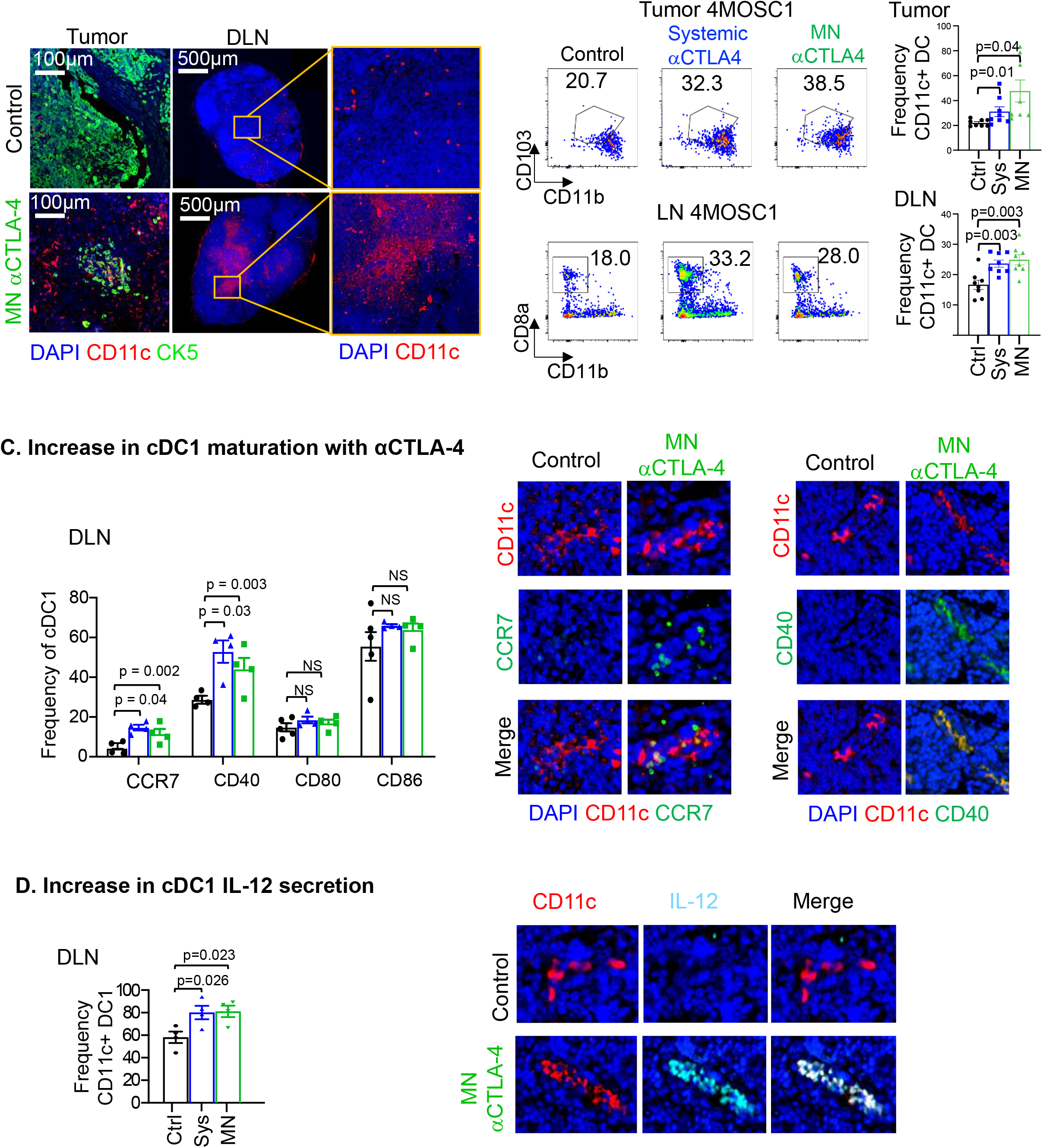
Recruitment of cDC1 DCs in tumor and draining LN by αCTLA-4. **A)** Immunofluorescence staining of CD11c (red) in tumors and cervical draining lymph nodes (DLN) highlights an increase in CD11c^+^ dendritic cells recruitment with αCTLA-4 treatment. Immunofluorescence staining of CK5 (green) show squamous cell character of the lesion and DAPI in blue. **B)** Flow cytometry analysis showing cDC1s increase intratumorally and in DLN with αCTLA-4 treatment comparing systemic and MN delivery strategies (n = 8 mice per group). **C)** Increase in cDC1 maturation markers with αCTLA-4 in the lymphatic compartment by flow cytometry (left, n = 3-5 mice per group) and representative IF staining (right). **D)** αCTLA-4-mediated increase of IL-12 secretion by cDC1s in the lymph node by flow cytometry (n = 3-5 mice per group) and representative IF stainings (right). Data are reported as mean ± SEM; two-sided Student’s *t*-test; the p-value is indicated where relevant when compared with the control-treated group; non-significant (ns).

### αCTLA-4 resistance in *Batf3* KO mice

The Batf3 transcription factor is critical for the development of cDC1s (18). To explore the role of Batf3-controlled cDC1s in the response to αCTLA-4, we transplanted our 4MOSC1 tumors into the tongues of *Batf3*-/- animals and found a general absence of intratumoral and lymph node cDC1s (**Supplemental Fig. 5**), and that αCTLA-4 had no antitumor activity in *Batf3* KO mice (**Fig. 3A and 3B**). Additionally, we observed an increase in the frequency of regional metastasis in tumor-bearing *Batf3*-/- animals compared to control, suggesting a role for cDC’s and likely their primed cytotoxic T cells in the control of regional disease spread in oral cavity cancer (**Fig. 3C**). Aligned with this possibility, MN delivery of αCTLA-4 resulted in significantly increased CD8^+^ T-cells infiltration, which was not observed in HNSCC tumors from *Batf3* KO animals by immunofluorescence (**Fig 3D**) and flow cytometry analysis (**Fig. 3E and 3F**). The latter also supported that the increase in the CD8+/CD4+ T-cell ratio caused by αCTLA-4 was dependent upon *Batf3* – regulated cDC1s.

**Fig. 3.**
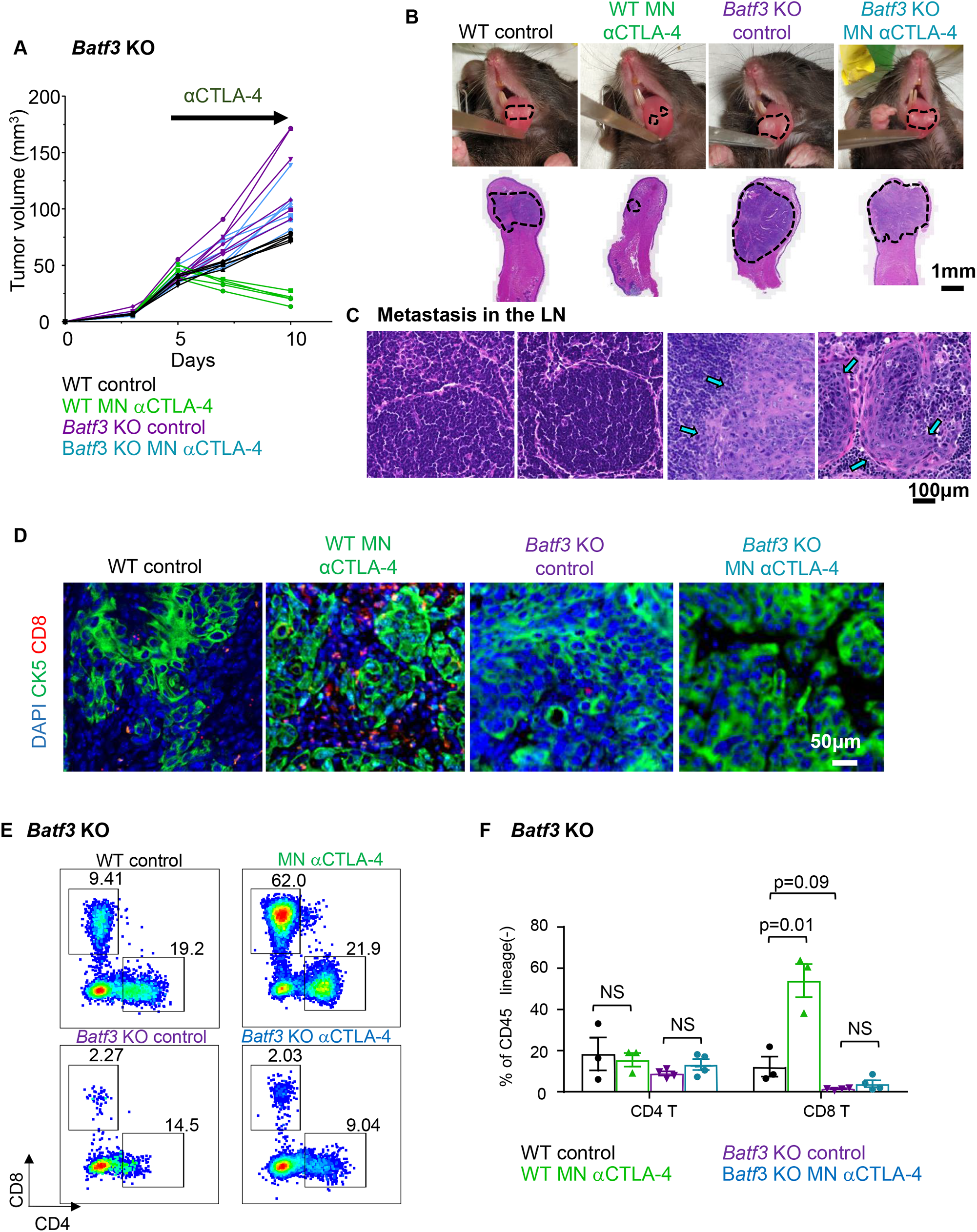
DCs are required for the αCTLA-4 response. **A)** Wild type and *Batf3* KO mice with 4MOSC1 tongue tumors were treated with negative PVP MN (control) or αCTLA-4 MN (0.1 mg) biweekly. Analyzed samples are related to the end of the study when the Batf3KO Ctrl mice had to be sacrificed. Individual growth curves of 4MOSC1 tumor-bearing mice are shown (n = 5-6 mice per group). **B)** Representative pictures of mice tongues and H&E-stained sections from panel **A**. Scale bars represent 1mm. **C)** Protective role of cDC1s in metastatic spread of 4MOSC1 to cervical lymph nodes. Representative H&E stain of cervical lymph nodes from mice in panel **A** are shown (n = 3 mice per group). Scale bars represent 100um. **D**) Immunofluorescence staining of CK5 and CD8 to show squamous cell character of the lesion (CK5^+^) and CD8 T-cell infiltration in mice from panel **A** (n = 3 per group) (CK5, green; CD8, red; DAPI, blue). **E)** Flow cytometry analysis of CD45^+^/CD11c-/Ly6G-/CD8^+^ or CD4^+^ T cell infiltration in WT or *Batf3* KO mice following αCTLA-4 treatment. A representative plot is shown (n = 3 mice per group) **F)** Quantification of tumor-infiltrating T cells from panel **E**. Data are reported as mean ± SEM; two-sided Student’s *t*-test; the p-value is indicated where relevant when compared with the control-treated group; NS, non-significant.

### Local MN αCTLA-4 delivery protects tumor-bearing mice from irAEs

Dose-limiting irAEs represent a considerable problem that impedes the clinical application of αCTLA-4 therapy with grade 3/4 toxic irAEs occurring after αCTLA-4 therapy in as many as 20-25% of cases (8). At baseline, wild type murine animal models are resistant to irAE development even with ICI delivered in combination, but Treg depletion in *FoxP3*-*DTR* (F*oxP3*^*DTR*^) genetically engineered mouse models (GEMMs) have been recently shown to lower the immune tolerance threshold and allow for irAEs to manifest after ICI therapy (20). In order to explore the relative toxicities of irAEs with systemic versus local αCTLA-4 therapy, we treated tumor-bearing wild type and *FoxP3*^*DTR*^ mice with systemic or local-MN αCTLA-4 in the presence or absence of acute diphtheria toxin treatment to achieve transient Treg depletion *in vivo* (20). We found that *FoxP3*^*DTR*^ animals recapitulate tumor growth kinetics observed in wild type hosts: all animals respond completely to αCTLA-4 systemic and MN treatment, and control-treated tumors grow progressively even after Treg depletion (**Fig. 4A**). Next, we performed a comprehensive analysis of irAEs in tumor bearing animals receiving therapy (**Fig. 4B-D** and **Supplemental Fig. 6**). While splenomegaly and blepharitis manifested in all animals receiving systemic αCTLA-4 treatment, local-MN αCTLA-4 treated mice were spared (**Fig 4B**). Moreover, histologic analysis of representative tissues (e.g., eyelid, skin, lung, liver, and colon) showed that delivery of αCTLA-4 local with our MN array spares tumor-bearing animals from the classic inflammatory reactions observed clinically and in preclinical models (12) (**Fig 4C** and **4D**), which was also reflected in weight gain in mice treated with MN delivered αCTLA-4 (**Supplemental Fig. 6**). Together, these results demonstrate that local MN therapy can achieve complete therapeutic response while protecting the host from development of irAEs.

**Fig. 4.**
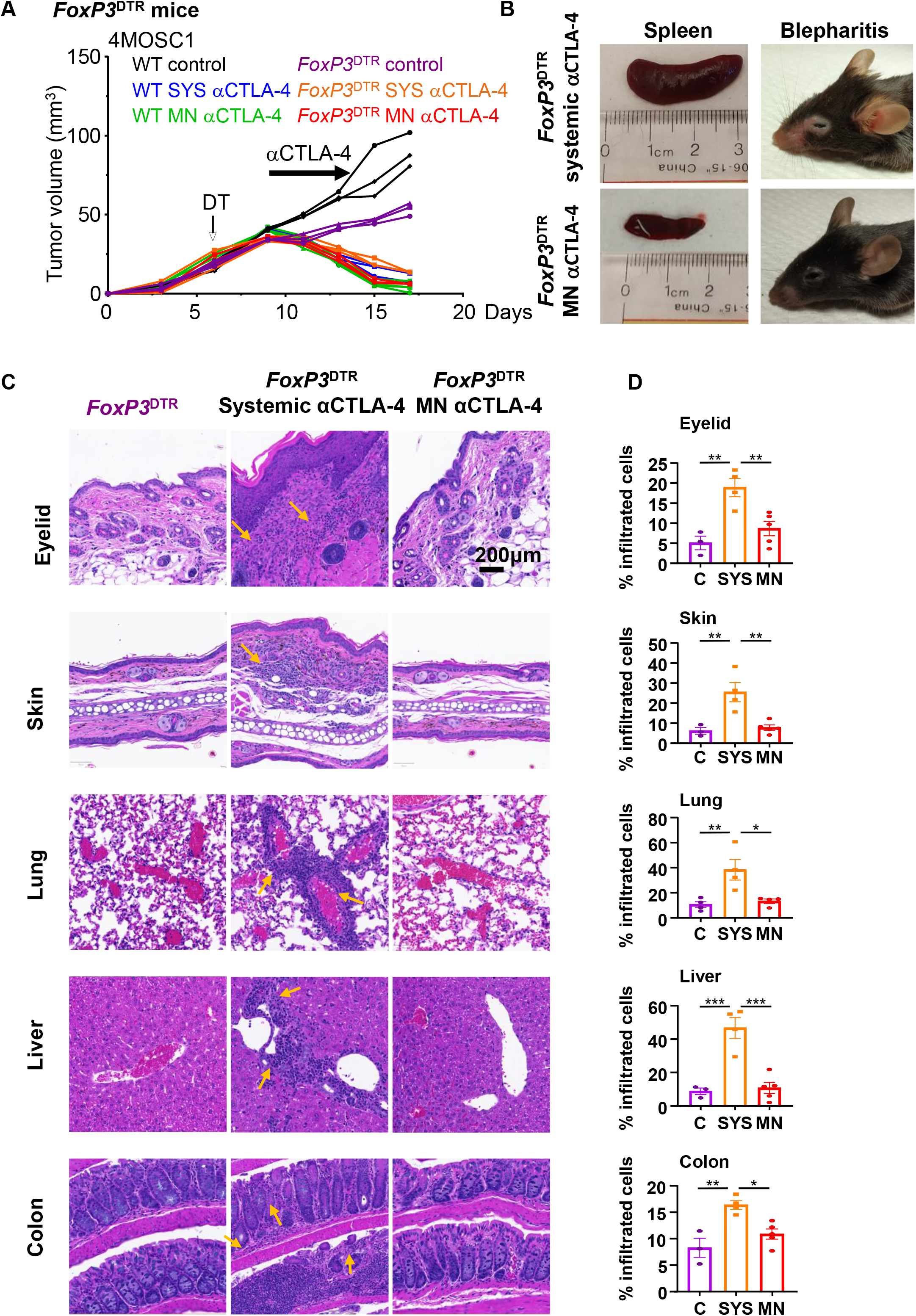
Local MN αCTLA-4 delivery protects tumor-bearing mice from irAEs. **A)** Wild type and *FoxP3*^*DTR*^ mice with 4MOSC1 tongue tumors were treated with negative PVP MN (control) or αCTLA-4 systemically (SYS; 5 mg/kg) or locally using MN (0.1mg per patch). *FoxP3*^*DTR*^ mice were treated by diphtheria toxin (DT, 500ng) to deplete FoxP3^+^ Tregs. Individual growth curves of 4MOSC1 tumor-bearing mice plotting primary tumor growth are shown (n = 3 mice per group). **B)** Representative pictures of spleen and eyes displaying splenomegaly and blepharitis, respectively, from mice in panel **A. C**) Representative H&E-stained sections of indicated organs from mice in panel **A**. Areas of inflammation are depicted by yellow arrows. **D**) Quantification of inflammatory immune cell infiltration in tissue sections from mice in panel **A** (n=3 mice per group). Data are reported as mean ± SEM; two-sided Student’s *t*-test; the p-value is indicated where relevant when compared with the control-treated group.

## Discussion

A core concept in cancer immunotherapy is that tumor cells, which would normally be recognized by T cells, have developed ways to evade the host immune system by exploiting mechanisms of peripheral tolerance (21,22). The use of novel and breakthrough immunotherapies, such as ICI, lower this threshold; and, in so doing, stimulate antitumor immunity to target and attack foci of primary and metastatic neoplastic disease (21,22). However, lowering the threshold of peripheral tolerance also induces off-target irAEs which ultimately temper the overall therapeutic benefit of ICI therapy. Therefore, novel therapeutic options and modalities for delivery are urgently needed to minimize ICI irAEs while preserving antitumor efficacy.

To address this clinical paradox, we have designed a novel MN system capable of delivering ICI with topical application and a targeted pharmacokinetic distribution limited to the primary tumor and draining lymphatic basins. In this regard, healthcare technology has seen significant advances in therapy as a result of implementing microfabrication tools towards the development of practical, precise and efficient alternatives for the delivery of pharmaceuticals (23). A variety of MN devices have been developed recently as a result of extensive preclinical testing by evaluating the safety and efficacy, e.g., vaccines and insulin delivery (24). MNs represent an attractive delivery system for local administration of anticancer therapeutics (25). Recently, dissolvable MNs were investigated for experimental melanoma treatment in mice by the delivery of anti-PD-1, achieving higher responses (∼40%) than when administered systemically (26). We now show that our MN delivery platform using αCTLA-4 achieves nearly complete tumor responses, and that remarkably, this IT αCTLA-4 administration protects the host from irAEs development.

Given the accumulation of αCTLA-4 antibody in the tumor and draining lymphatic compartments along with the requirement for CD8 T-cells to mediate tumor rejection, we explored upstream to interrogate the intervening cellular immune mechanism. We found that Batf3-cDC1s are strictly required for *in vivo* αCTLA-4 tumor responses. Stemming from their unique ability to cross-present antigen and bridge CD4 and CD8 T-cell priming (19), Batf3-cDC1s are now appreciated as the critical upstream immune effector in priming antitumor immune responses (18). The unique responsiveness of the 4MOSC1 model to monotherapeutic αCTLA-4 afforded the opportunity to extend the wealth of knowledge regarding Batf3-cDC1s to the mechanism of action of αCTLA-4 ICI. Our findings document a strict requirement for host Batf3 downstream from αCTLA-4 ICI, thus supporting a role for classical cDC1-CD8 T-cell priming for αCTLA-4 efficacy. This novel observation offers insight into possible multimodal co-targeting strategies. y virtue of it’s localized mechanism of action, the MN delivery system is an ideal approach by which to explore various combinations of IO therapies, such as those that include tandem local and systemic treatment, such as local MN αCTLA-4 delivery with systemic αPD-1/αPD-L1 treatment.

Whether effective anti-tumor response to MN αCTLA-4 requires modulation of other immune cell types, such as the myeloid cell compartment, requires further investigation. On the other hand, as ICI therapies become increasingly standard for cancer therapy, amelioration of irAEs becomes an increasingly important goal. In light of this, many groups are developing elegant and specific preclinical models to examine irAEs (20,27). Here, we employed the F*oxP3*^*DTR*^ GEMMs to abrogate the immune tolerance threshold and permit irAEs to manifest after ICI therapy. An alternative and recently characterized model – the dextran sulfate sodium colitis murine model – engenders a graft-versus-host environment *in vivo* by adoptively transferring human peripheral blood mononuclear cells and, subsequently, human colon cancer cells. This model faithfully reproduced ICI-associated colitis (27), and hence can be also used in future studies to interrogate the translational applications of our MN delivery system to reduce the development of irAEs. In this regard, our study is largely focused on adverse events resulting from αCTLA-4, but the emerging results can now be readily extended to the use of MN to deliver other ICIs as single agents and as part of combination immunotherapies.

Taken together, our findings demonstrate that targeted, local IT delivery of αCTLA-4 ICI will initiate a robust and durable antitumor response that is dependent on cDC1 and CD8+ T-cells, while significantly limiting irAEs. In this regard, the use of an innovative MN patch may afford a feasible and effective local-delivery platform for IO agents that can be quickly translated into the clinical setting.

## Acknowledgements

This project was supported by grants from National Cancer Institute (R01CA247551), and National Institute of Dental and Craniofacial Research (NIH/NIDCR, U01DE028227) and NIH grant S10OD021831 to Dr. Z. Mikulski. M.A.L.-R. and F.S. acknowledge the UC MEXUS-CONACYT Doctoral Fellowships. This work was partially supported by Salk Cancer center pilot award CA014195 to Dr. D.P. Hollern. Some schematics and figures were created with biorender.com. We thank Drs. Scott M. Lippman, Joseph A. Califano, Jesse Qualliotine, Jayanth S. Shankara Narayanan, and the Staff of La Jolla Institute Microscopy Core and Flowcytometry core Facilities for insightful suggestions.

