## Supplemental materials and methods for "Microneedle-mediated intratumoral delivery of anti-CTLA-4 promotes cDC1-dependent eradication of oral squamous cell carcinoma with limited irAEs"

### Supplemental Information

### Supplemental materials and methods

#### αCTLA-4 MN patch characterization

The fluorescent microscopy images of MN patches were obtained with the use of an EVOS FL microscope precisely coupled with 2X and 4X objectives, as well as the corresponding fluorescence filter, for red light excitation (RFP) of Alexa Fluor 555. The scanning electron microscopy (SEM) images were captured by the employment of a FEI Quanta 250 ESEM instrument (Hillsboro, Oregon, USA), using an acceleration voltage of 3 kV. The MN patch was previously sputtered with Iridium for better visualization.

#### αCTLA-4 MN patch dissolution test

The MN patches were sliced in 3x1 MN array pieces and placed horizontally against a clear glass slide; subsequently, the dissolution of these patches was performed by the addition of PBS pH 7.4 at an inverted optical microscope (Nikon Eclipse Instrument Inc. Ti-S/L100) coupled with a 4x microscope objective, a Hamamatsu digital camera C11440, and a NIS Elements AR 3.2 software.

#### Released IgG Alexa Fluor-555 detection

The MN patches were set to penetrate a synthetic mimicking phantom tissue. Briefly, 2% agarose was fabricated in distilled water and casted into custom made negative well EcoFlex molds of 3mm in thickness. The resulting circular phantom tissues were submerged in Phosphate Buffer Saline (PBS) pH 7.4 and placed in a sealed container until use. The MN patches pierced the mimicking tissue barrier at several time set points (1,2,3,5,10,20 and 30 min) at 37.5 °C; after the MN application, the remaining material of the patch was removed, and dissolved in 800µL of PBS pH 7.4. The absorbance was subsequently measured from 400 to 700 nm using a UV-2450 Shimadzu spectrophotometer and the cumulative release was plotted vs time.

### Supplementary figure legends

**Supplemental Fig. 1.** Microneedle fabrication and characterization. **a)** Schematic of a dissolvable microneedle application on 4MOSC1 tumor of mouse for the delivery and release of an immune oncology agent (αCTLA-4) to treat HNSCC and digital photograph of a dye supplemented microneedle application into a synthetic dye supplemented hydrogel mouse tongue. Scale bar, 3mm. **b)** Fabrication steps of the microneedle patch: PDMS micromolding over master microneedle, PDMS negative MN mold released, polymer and payload loading, polymer air-drying, adhesive application and release. **c)** Digital photograph showing a dissolvable patch comprised of 64 microneedles, and a scanning electron micrograph of the microneedle array. Scale bar, 400µm. **d)** Fluorescent microscopy image of 3 microneedle tips loaded with Alexa Fluor-555, scale bar, 500µm respectively. **e)** Microscopy time-frame images of a single microneedle tip clearly showing polymer dissolution and further payload release. **f)** Cumulative release kinetics curve of a model payload (Rh-6G) released from passive microneedles.

**Supplemental Fig. 2.** C57Bl/6 mice were implanted with $1 \times 10^6$ 4MOSC1 cells into the tongue. After the tumors reached ~30 $mm^3$, mice were treated with isotype control or different doses of αCTLA-4 for systemic administration. The average tumor volume of each group is shown (n = 5 mice per group, data are represented as mean ± SEM).

**Supplemental Fig. 3. a)** Gating strategy of intratumoral and lymphoid cDC1s. Dendritic cells were characterized as Thy1.2-, CD11c+, and MHCII+. From there, intratumoral cDC1s from the tumor were characterized as CD11c+, CD103-, and CD11b-, and lymphoid cDC1s from the DLN were characterized as CD11c+, CD8a+, and CD11b-. **b)** Representative flow cytometry plots showing characterization of maturation markers from lymphoid cDC1s. The frequencies of CD40, CCR7, CD80, and CD86 out of lymphoid cDC1s were calculated following treatment with MN or systemic αCTLA-4.

**Supplemental Fig. 4.** Quantification of CD8 T cells in the LN following anti-CTLA-4 treatment by flow cytometry. The absolute number of CD8 T cells were calculated in each DLN following treatment with MN or systemic αCTLA-4. Data are reported as mean ± SEM; two-sided Student's *t*-test; the *p*-value is indicated where relevant when compared with the control-treated group; non-significant (ns).

**Supplemental Fig. 5.** Representative flow cytometry plot showing intratumoral and lymphoid cDC1 DCs in *Batf3* KO mice compared to WT mice. The gating strategy for cDC1s are described in Supplemental Fig. 3.

**Supplemental Fig. 6.** *FoxP3*$^{DTR}$ or WT mice were treated with MN or systemic $\alpha$CTLA-4. After 6 treatments, mice were weighed. *, $p<0.05$.
