## Supplemental figures for "Microneedle-mediated intratumoral delivery of anti-CTLA-4 promotes cDC1-dependent eradication of oral squamous cell carcinoma with limited irAEs"

**Supplemental Fig. 1 – Microneedle patches design**

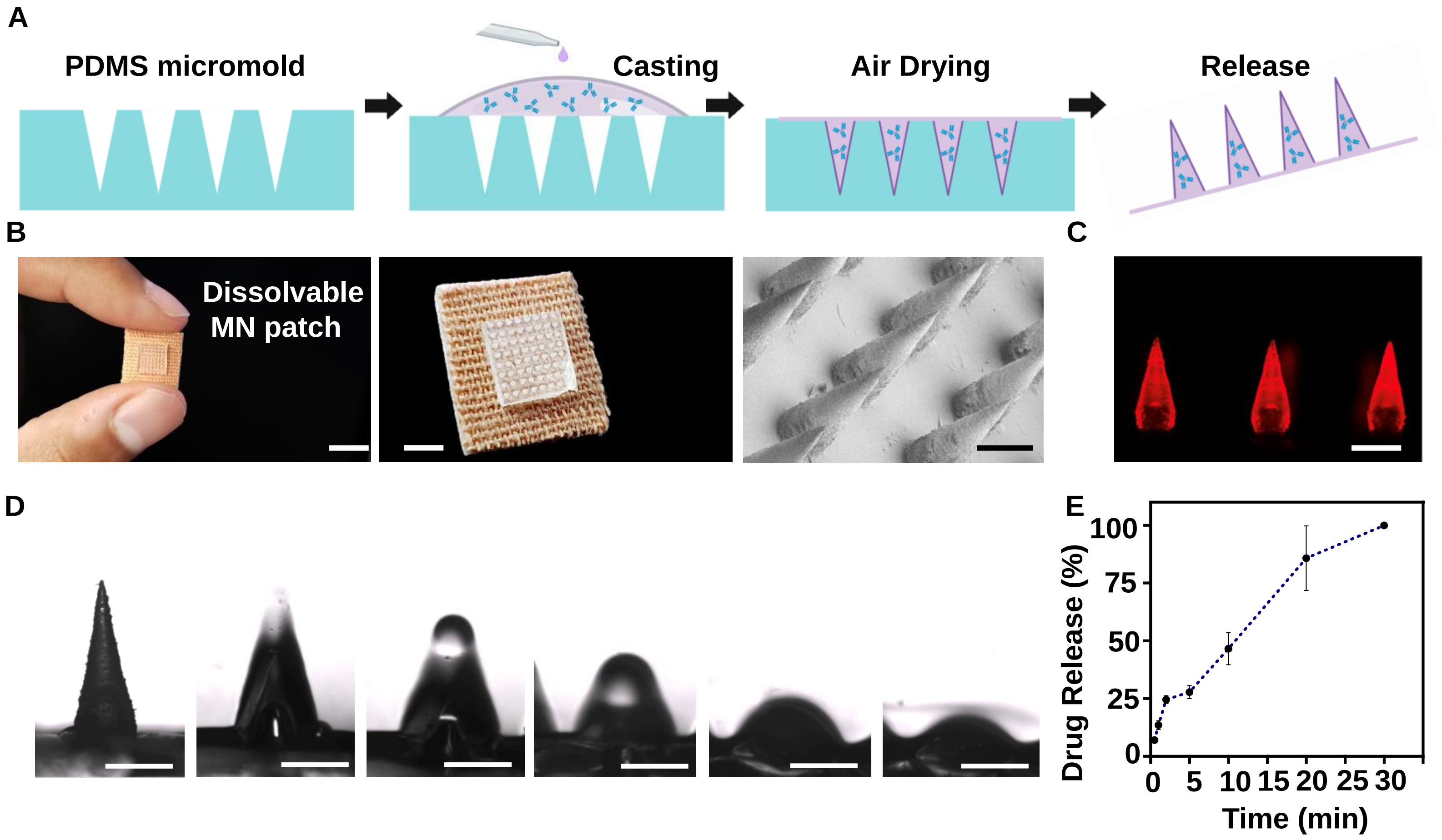

**Supplemental Fig. 2 – Dose escalation of CTLA4 to identify the concentration to use in the MN patch**

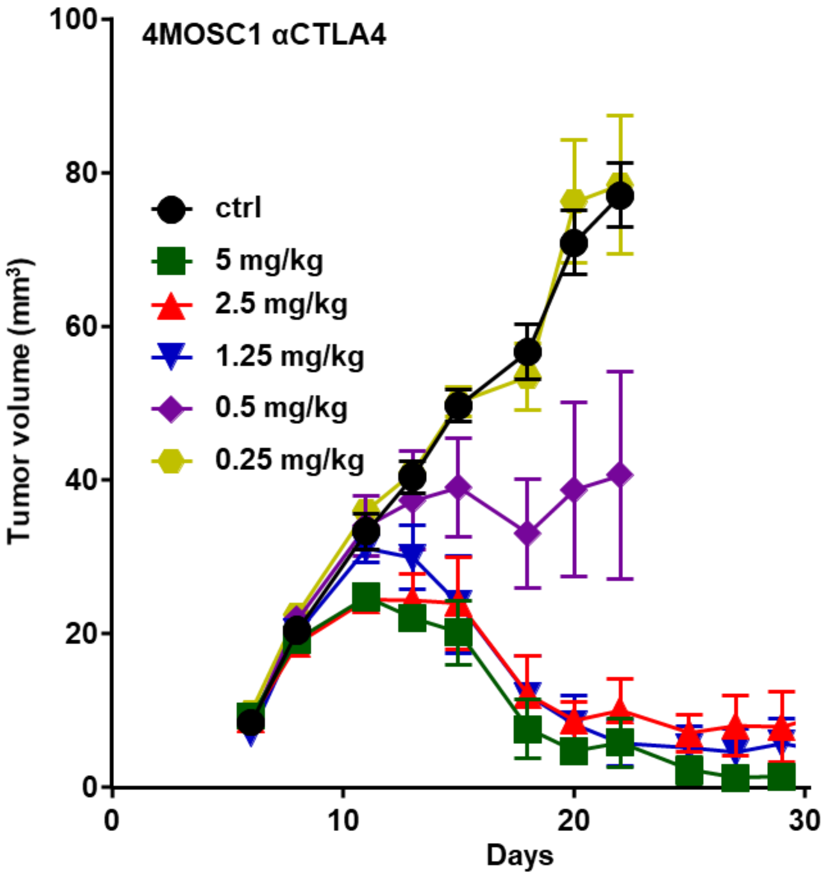

**Supplemental Fig 3 - cDC1 phenotype and maturation markers gating strategies**

Intratumoral – CD11c$^{+}$CD103$^{+}$CD11b$^{-}$
Lymphoid – CD11c$^{+}$CD8a$^{+}$CD11b$^{-}$

A

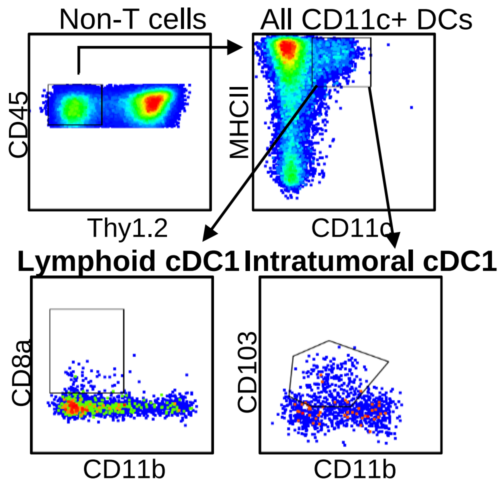


B

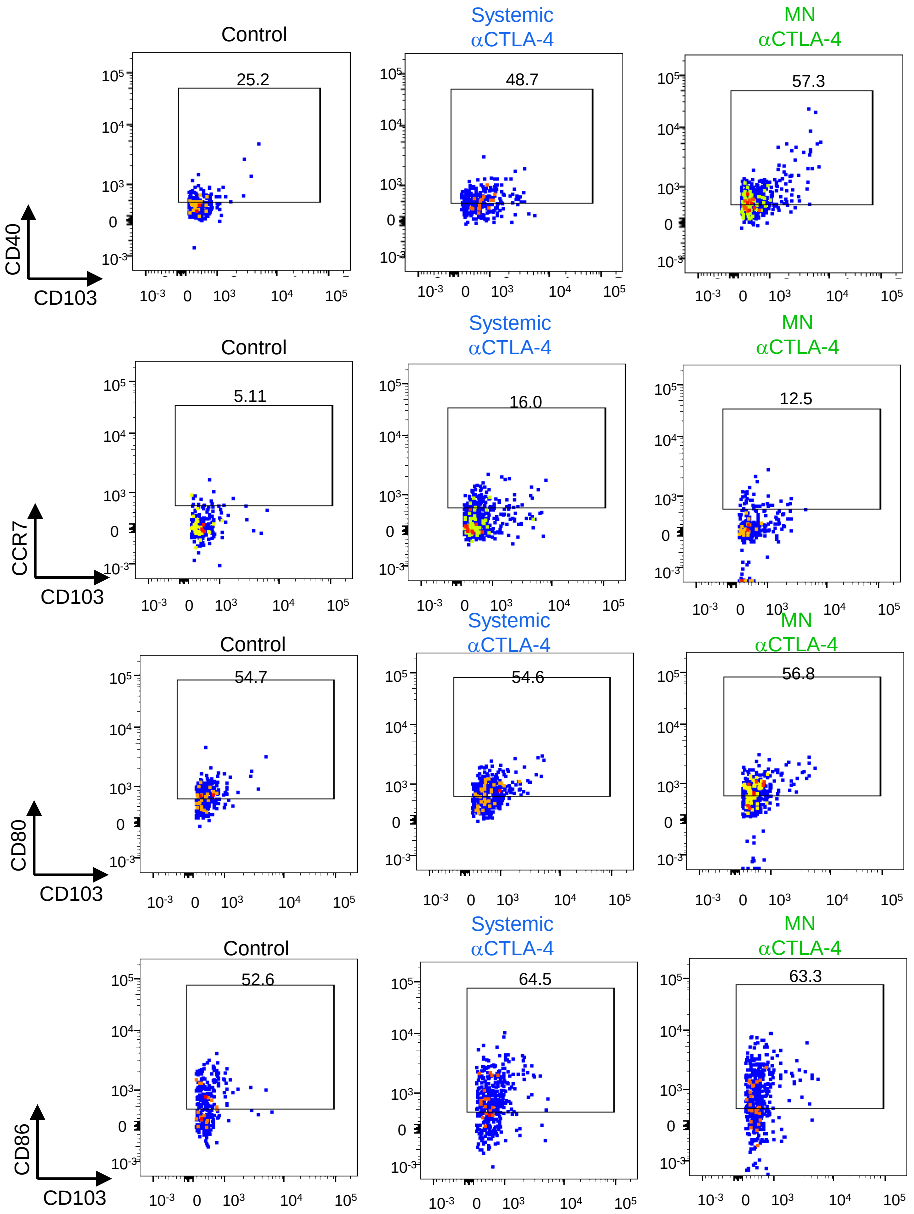

**Supplemental Fig. 4 – Quantification of CD8 T cells in the LN following anti-CTLA-4 treatment**

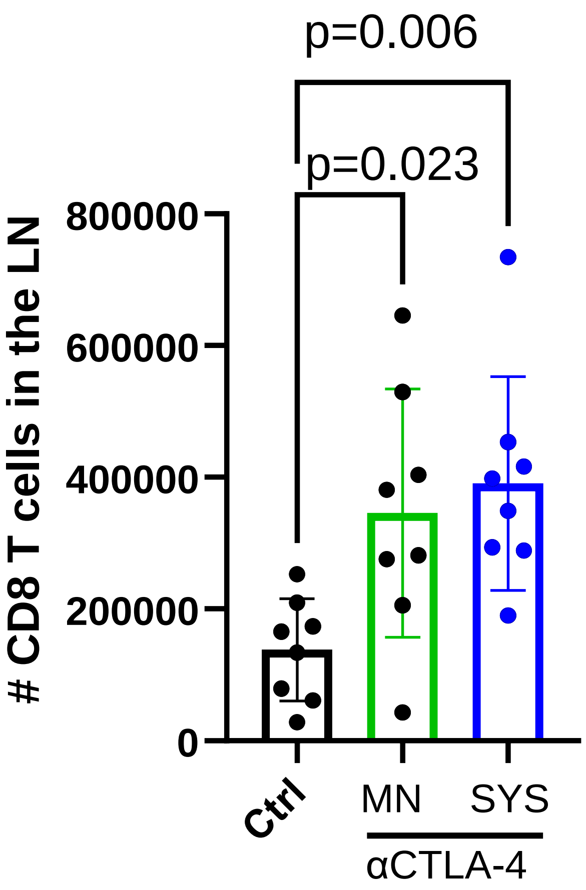

**Supplemental Fig. 5 – Intratumoral and lymphoid cDC1 DCs in *Batf3* KO mice**

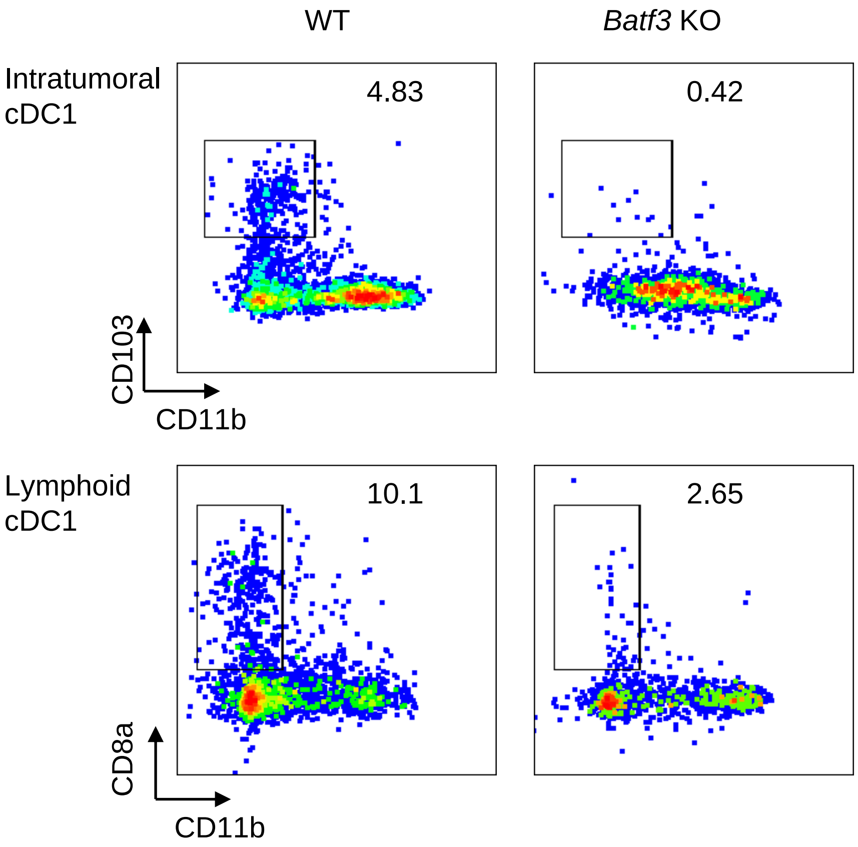

**Supplemental Fig. 6 – Weight of the *FoxP3*$^{DTR}$ mice during different treatments**

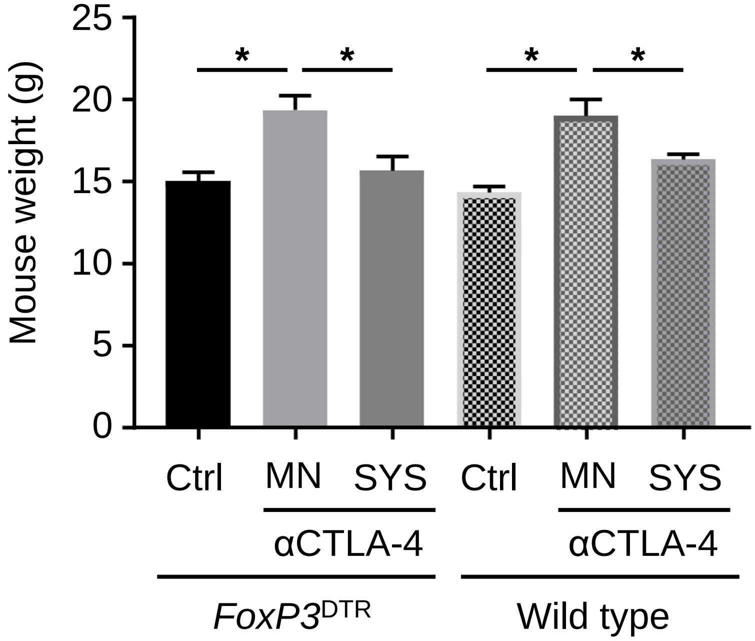
